# Solvent based fractional biosynthesis, phytochemical analysis and biological activity of silver nanoparticles obtained from extract of *Salvia moorcroftiana*

**DOI:** 10.1101/2023.05.30.542991

**Authors:** Maham Khan, Tariq Khan, Muhammad Aasim, Tauqir A. Sherazi, Shahid Wahab, Muhammad Zahoor

## Abstract

Multi-drug resistant bacteria sometimes known as “superbugs” developed through overuse and misuse of antibiotics are determined to be sensitive to small concentrations of silver nanoparticles. Various methods and sources are under investigation for the safe and efficient synthesis of silver nanoparticles having effective antibacterial activity even at low concentrations. We used a medicinal plant named *Salvia moorcroftiana* to extract phytochemicals with antibacterial, antioxidant, and reducing properties. Three types of solvents; from polar to nonpolar, i.e., water, dimethyl sulfoxide (DMSO), and hexane, were used to extract the plant as a whole and as well as in fractions. The biosynthesized silver nanoparticles in all extracts (except hexane-based extract) were spherical, smaller than 20 nm, polydispersed (PDI ranging between 0.2 and 0.5), and stable with repulsive force of action (average zeta value = −18.55±1.17). The tested bacterial strains i.e., *Klebsiella pneumoniae, Pseudomonas aeruginosa, Staphylococcus aureus, and Enterococcus faecali*s were found to be sensitive to even small concentrations of AgNPs, especially *P. aeruginosa.* The antibacterial effect of these AgNPs was associated with their ability to generate reactive oxygen species. DMSO (in fraction) could efficiently extract antibacterial phytochemicals and showed activity against MDR bacteria (inhibition zone = 11-12 mm). Thus, the antibacterial activity of fractionated DMSO extract was comparable to that of AgNPs because it contained phytochemicals having solid antibacterial potential. Furthermore, AgNPs synthesized from this extract owned superior antibacterial activity. However, whole aqueous extract based AgNPs MIC was least (7-32 µg/mL) as compared to others.

## 1. Introduction

Since the discovery of penicillin, the population across the globe has relied on antibiotics to fight microbial infections. Regrettably, microbes have found a way to resist the action of antibiotics that were once effective due to their misuse, combined use, and over-prescription (1). The development of multi-drug resistant microbes has now become a universal threat with severe results in administering infectious illnesses caused by pathogens (2). The need to overcome this problem has been well recognized and highly investigated. Researchers are coming up with solutions to combat antibiotic resistance, among which nanotechnology is leading the race. Nanoparticles have emerged as unique particles with size-dependent physicochemical characteristics that are proven to have antibacterial potential (3). Silver nanoparticles (AgNPs) have attracted attention as they hold great promise in combating the challenge of antimicrobial resistance due to their extraordinary, broad-spectrum, and strong antimicrobial properties (4).

For the efficient synthesis of nanoparticles, green methods employing microbial cells, plant extract, and natural polymers have been developed. These methods are cost-effective, environment-friendly, and energy saving (5–7). Among the available biosynthetic methods, the synthesis of nanoparticles using plant extract is the most favorable. It is a safe and green method that can be easily scaled up for producing a large amount of nanoparticles (8). Plant metabolites actively participate in the synthesis of nanoparticles by reducing and capping metallic cations. These phytochemicals are known to possess good antimicrobial potential and have been attributed to the increased antimicrobial effect of nanoparticles capped by them in the process of biological synthesis through plant extracts. Alkaloids, acetylenes, coumarins, flavonoids and isoflavonoids, iridoids, lignans, macrolides, phenolics (other than flavonoids and lignans), polypeptides, quinones, steroidal saponins, terpenoids, and xanthones have been identified to possess strong effect against resistant bacteria (Saleem et al 2010). Bearing in mind the antibacterial properties of these compounds, which have been used for centuries, are a source of new therapeutic agents.

The extraction of phytochemicals depends on the type of extraction technique and nature of solvent because the different polarity and chemical characteristics may or may not allow the dissolution of phytochemicals in particular solvents. Thus, various solvents have different affinity for phytochemicals from plants (9). Phenolic acids, flavonoids, tannins, and alkaloids etc., can be extracted in polar or less polar solvents (10). In comparison, components of essential oils and tocopherol can be obtained through extraction in nonpolar solvents (11). Thus, solvent based plant extracts can help us to reach the specific group of plant compounds that is responsible for the synthesis of nanoparticles with high bactericidal activity, stability, and dispersity.

*Salvia moorcroftiana,* commonly known as “kallijari” in Pakistan is a herbaceous plant known to have medicinal properties to relieve pain, fever, and inflammation (12). The plant contains valuable phytochemical contents like essential oils, flavonoids and polyphenols, tannins, terpenoids, phytosterols, carbohydrates, etc. (13). The polyphenols and flavonoids are natural antioxidants and possess other pharmacological activities such as anticancer, anti-inflammatory, antianxiety, and antimicrobial (14). The targeted extraction of plant metabolites possessing antibacterial properties can be obtained by using suitable solvents (in terms of polarity) for plant extraction (15). The metabolites in *S. moorcroftiana* possess the potential of AgNPs synthesis with varying and enhanced antibacterial activity. To the best of our knowledge, no study is available that attempts to synthesize and study AgNPs synthesized through this plant using different solvents. Thus, this research study focuses on the antibacterial activity and bioreduction potential of *S. moorcroftiana* extracts based on highly polar (Water), less polar (Dimethyl Sulfoxide), and nonpolar (n-hexane) solvents. The purpose of our current study was to determine the effect of polarity of solvents and the extraction method on the extraction of biologically important phytochemicals of *S. moorcroftiana* and the synthesis of AgNPs (Fig 1). This study further aims to comparatively analyze AgNPs synthesized from the extracts of *S. moorcroftiana* in different solvents, in terms of stability, morphology, functional groups, surface charge, and activity against multi-drug resistant bacteria.

**Figure 1:**
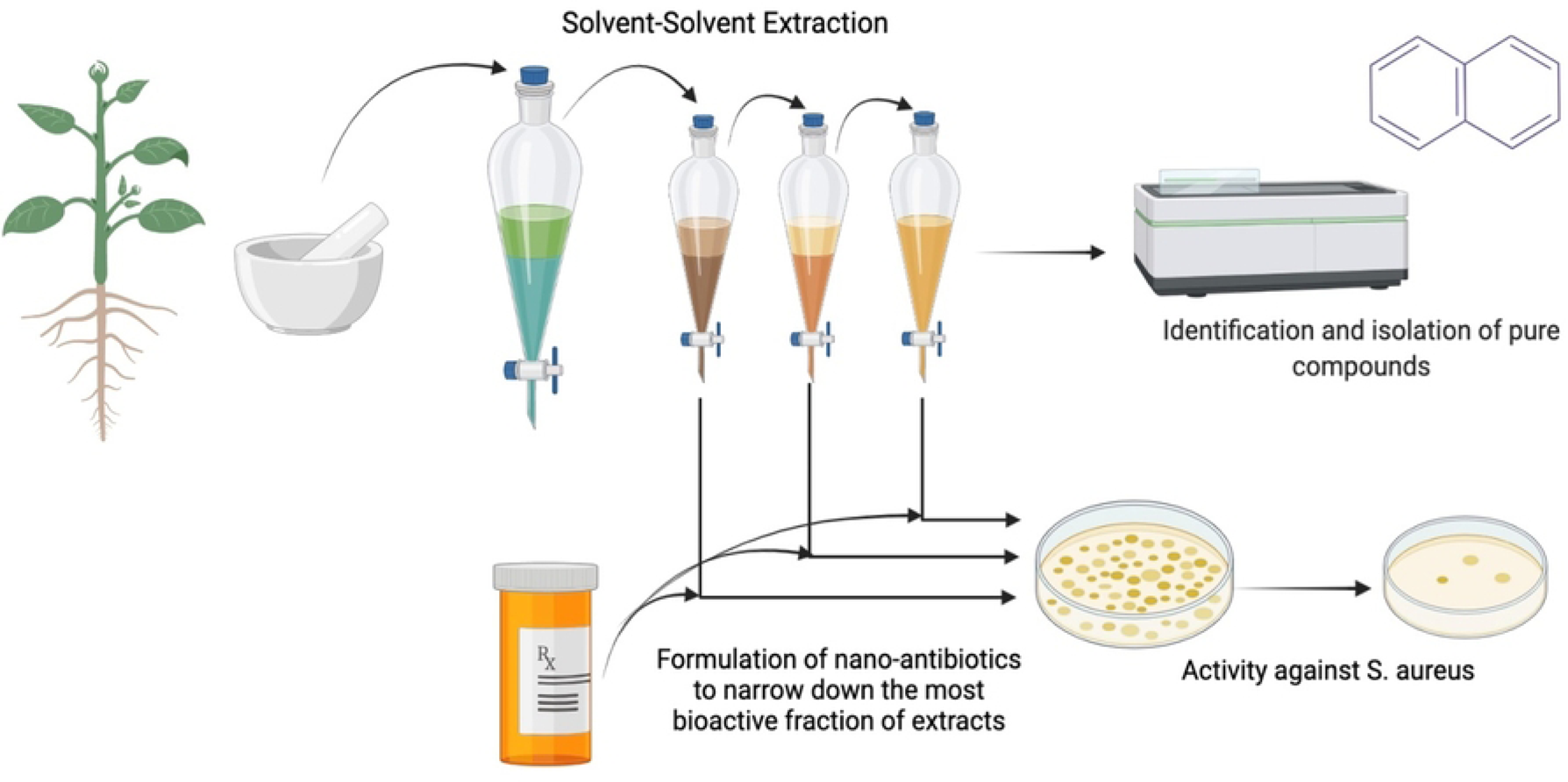
Solvent-based extraction of antimicrobial phytochemicals

## 2. Materials and Methods

### 2.1 Plant material and preparation of extract

Fresh *Salvia moorcroftiana* leaves were collected from district Swat, Khyber Pakhtunkhwa, Pakistan. The plant was shade dried and grinded into fine powder for extraction. Three types of extracts were prepared in different solvents ranging from polar to nonpolar: aqueous extract (Aq-extract), DMSO based extract (D-extract), and hexane-based extract (H-extract).

#### 2.1.1 Preparation of aqueous extract

Aqueous extract of *S. moorcroftiana* was prepared by mixing 5 g of plant powder with 100 mL distilled water. The mixture was boiled for 10mins and cooled at room temperature. The solution was filtered through Whatman (2.5 µ) filter paper to completely remove the plant remains. The resultant extract was stored at 4°C for further use.

#### 2.1.2 Dimethyl sulfoxide based extraction

DMSO based extract was prepared by mixing 2 g of plant powder with 50 ml DMSO. The mixture was sonicated for 30mins. Whatman (2.5 µ) filter paper was used to filter the mixture to remove plant debris. The dark green viscous plant extract in DMSO was obtained.

#### 2.1.3 Hexane based extraction

Hexane based extract was prepared by mixing 5g of plant powder with 100ml hexane. The mixture was heated in water bath at 60°C for 20mins and then agitated in a shaker for 30mins. The mixture was filtered through Whatman filter paper. The thin hexane based extract with light green color was obtained.

#### 2.1.4 Solvent-solvent fractional extraction of *Salvia moorcroftiana*

To separate phytochemicals based on their relative solubility liquid-liquid extraction was used. 5g of plant powder was mixed in 100 mL hexane which was then heated at 60°C for 20 mins, agitated for 30 mins in a shaker and then filtered through Whatman filter to extract the nonpolar components of plant. The filtrate was stored at 4°C and the plant debris was used for further extraction. To the remaining plant debris 100 ml DMSO was added and sonicated for 30mins. The mixture was filtered through Whatman filter. The filtrate termed DMSO*-extract was stored at 4°C. To the remaining plant debris 100 ml distilled water was added and boiled for 10mins. Subsequently, the mixture was filtered and Aq*-extract was obtained that was stored for further use.

### 2.2 Biosynthesis of Silver nanoparticles

AgNPs were synthesized in all fractions of extracts of *S. moorcroftiana.* For the synthesis, both aqueous extracts i.e. obtained directly from intact plant material (Aq-extract) and through solvent-solvent extraction (Aq*-extracts) were mixed with 4 mM solution of Silver nitrate (AgNO_3_) in clean flasks in a specific ratio (1:2). This means to 100 mL aqueous plant extract 200 mL AgNO_3_ solution was added. The solution was incubated in daylight for 3 hrs. Similarly, DMSO based extracts obtained directly from intact plant powder (D-extract) and through solvent-solvent extraction (D*-extracts) were mixed with AgNO_3_ solution in 1:6. This means that 20mL DMSO based extract was mixed with 120 mL AgNO_3_ solution. The solution was incubated in daylight for 4 hrs. The hexane based extract (30 mL) was dried in oven at 50°C for about 6 hrs. After this an oily semi-dried extract was obtained. Methanol, due to its known ability to extract broad range of phytochemicals was used to dissolve compounds from dried hexane extract. 20 mL methanol was added to the dried hexane extract and the mixture was vortexed for 15 mins followed by sonication for 5 mins. The resultant methanolic extract was filtered (Whatman filter) to remove clumps. Finally, 30 mL of 4mM AgNO_3_ solution (synthesized in methanol) was added to 15 mL methanolic extract and incubated for 24 hrs.

The synthesis of AgNPs was observed in each of the extract by a change in color. Synthesis in aqueous extract changed the color of solution from light brown to reddish brown; in DMSO extract the color changed from light green to dark brown, and in hexane extract the color changed from lime to light grey. The AgNPs synthesized in all the extracts were obtained through centrifugation at 13000 rpm for 10 mins. The pellet (nanoparticles) was dried in oven at 50°C for 24 hrs, collected, and stored.

### 2.3 Characterization and confirmation of silver nanoparticles

Characterization of all types of AgNPs was carried out. These nanoparticles included AgNPs synthesized in aqueous extract of *S. moorcroftiana* (Aq-AgNPs), in aqueous fraction (Aq*-AgNPs), in DMSO based extract (D-AgNPs), and DMSO based fraction (D*-AgNPs).

#### 2.3.1 UV-Visible spectroscopy

The biosynthesis of AgNPs was confirmed by measuring absorbance using UV-Visible spectroscopy (BKS360 BIOBASE Spectrophotometer). 1 mL of each sample of reaction mixture (after reaction of AgNO_3_ and plant extract) was scanned in a spectrophotometer with the wavelength ranging from 300-600 nm. Each of the respective plant extracts were taken as control. The solvents used for the synthesis of AgNPs were taken as reference like for AgNPs synthesized in aqueous extract water was used as a reference.

#### 2.3.2 Fourier Transform Infrared Spectroscopy

FTIR characterize the biomolecules involved in the formation and stabilization of AgNPs. To indicate the phytochemicals involved in the capping of AgNPs in each of the extract of *S. moorcroftiana* including aqueous extracts i.e., Aq-extract, Aq*-extract D-extract, and D*-extract. All types of AgNPs (2 mg) in their dried form were grinded with potassium bromide (KBr) pellets. For appropriate results verification, scanning was done at a 1:100 ratio followed by recording a spectrum at a wavelength of 4500 to 500 cm^-1^. The different modes of vibrations of functional groups were identified and associated to the belonging biomolecules.

#### 2.3.3 Transmission Electron Microscopy

The size and shape of biosynthesized AgNPs was calculated and visualized through TEM. 2 µL of AgNPs suspension were dropped onto 400 mesh carbon-coated copper grid. The suspension was dried under a lamp in ambient conditions to form a thin layer. Afterward, the samples were subjected to the TEM wherein the interaction of transmitted electrons and the AgNPs formed images that were recorded. The average size of nanoparticles was determined by measuring the diameter of more than 50 particles on 100 nm TEM image.

#### 2.3.4 Dynamic light scattering of Silver nanoparticles

Dynamic Light Scattering (DLS) is a method used for the determination of size distribution of the nanoparticles by measuring radiation scattering intensity based on the Brownian motion of particles. DLS is a non-destructive technique used to measure size distribution and stability of silver nanoparticles in aqueous or physiological solutions. DLS uses light scattering to determine the average hydrodynamic diameter of the sample. A beam of light is put on dispersion of nanoparticles, which scatter light to the detector. The polydispersity index (PDI) is depicted as the size distribution range of nanoparticles and their stability and uniformity. From this measurement, the mean size of particles inside the sample is obtained along with the correlation between the numbers of particles of a particular size versus the size of the nanoparticles

#### 2.3.5 Zeta potential determination for stability assessment

Zeta potential analysis of AgNPs synthesized in each aqueous and DMSO based extract was performed to determine their suspension stability. 3 mg AgNPs/5 mL dH_2_O was prepared and sonicated for 30 mins (Power-Sonic 405). The surface charge and hydrodynamic diameter of nanoparticles samples were analyzed using a zeta sizer (Nano-ZS ZEN 3600, Malvern) with a temperature equilibration time of 1 minute at 25°C.

### 2.4 Antibacterial activity

#### 2.4.1 Microbial strains

The antibacterial activity of each type of extract (Aq-extract, Aq*-extract, D-extract, D*-extract, and H-exract) and AgNPs (Aq-AgNPs, Aq*-AgNPs, D-AgNPs, D*-AgNPs, and H-AgNPs) was performed. The tested microorganisms include Gram negative *Klebsiella pneumoniae* and *Pseudomonas aeruginosa,* and Gram positive *Enterococcus faecalis* and *Staphylococcus aureus*.

#### 2.4.2 Disk diffusion method

The Kirby-Bauer disk diffusion assay was performed to check the antibacterial activity of nanoparticles and *S. moorcroftiana* extracts. Each of the bacterial strain was streaked on nutrient agar solidified in sterile petri-dishes. After 24 hrs of incubation at 37°C, 3-5 colonies were picked using sterile metal loop and were inoculated in fresh broth medium contained in sterile glass tube with screw caps. The broth inoculated with bacteria was vortexed followed by 20 mins incubation. The broth suspension was compared with 0.5 MacFarland standard and the turbidity was adjusted accordingly. After adjusting turbidity of broth suspension according to MacFarland standard indicates 1×10^8^ CFU/mL of bacteria in broth. The bacteria were uniformly spread on nutrient agar in sterile petri-dishes using a sterile cotton swab dipped in broth suspension culture. The empty disks prepared from Whatman filter paper were placed on the plate (containing agar swabbed with bacteria) using autoclaved forceps. 8 µL of 2 mg/ml of each AgNPs and extract sample was added to each respective disc through sterile micropipette tips. The petri-dishes were sealed with parafilm and incubated for 24 hrs at 30°C. The inhibition zones were recorded after 24 hrs of incubation.

#### 2.4.3 Micro-dilution assay for determination of minimum inhibitory concentration

Micro-dilution assay for each type of nanoparticles was performed to determine the minimum inhibitory concentration. The pure culture of each bacterial strain was obtained by streaking the bacteria which was incubated for 24 hrs at 37°C. The colony suspension method was followed to prepare bacterial broth culture. 3-5 colonies of bacteria were picked with the help of sterile metal loop and were inoculated in fresh broth medium. The bacterial broth culture was compared with 0.5 MacFarland standard by comparing the optical density (OD) at 625 nm. The absorbance was observed in the range of 0.08-0.13 which is the desired value and indicates approximately 1×10^8^ CFU/ mL. 8 mg/mL stock solution of Aq-AgNPs, Aq*-AgNPs, D-AgNPs, D*-AgNPs, and 4 mg/mL H-AgNPs (stock solution) was prepared. The stock solutions were twofold diluted up to 10^th^ well. The concentrations of Aq-AgNPs, Aq*-AgNPs, D-AgNPs, and D*-AgNPs started from 4 mg/ml and ended at 7.8 µg/mL. However, H-AgNPs the concentrations started from 2 mg/ml and ended at 3.9 µg/mL.

The dilutions were added to their respective wells of 96-well microtitre plate. Separate plate was used for each of the bacteria. 50 µL of bacterial broth culture was added from 1^st^ to 11^th^ (growth control) column. 50 µL of broth (only) was added to the 11^th^ column and 100 µL to the 12^th^ (sterility control). The plate was incubated for 20 hrs at 37°C in a shaking incubator. The MIC was observed visually by determining the concentration at which AgNPs inhibited the bacterial growth resulting in a clear well.

#### 2.4.4 Mechanism of bacterial inhibition by the biosynthesized AgNPs

The oxidant sensitive probe 2’, 7’-dichlorodihydroluorescein diacetate (H_2_DCFDA) was used to determine the intracellular levels of ROS in cells treated with different concentrations of AgNPs (0.5-250 µg/mL). Bacterial cells were grown in LB medium until OD_600_ reaches 0.5. The bacterial cultures were centrifuged to pellet the cells. The cells were washed with 10 mM potassium phosphate buffer (pH 7.0) through vortexing and centrifugation. After washing, the cells were suspended in the same buffer and disrupted by sonication. 10 mM H2DCFDA (dissolved in dimethyl sulfoxide) was added at a ratio of 1∶2000 (2 microliter in 4 ml buffer), followed by shaking for 30 min at 37°C.the bacterial cells were again pelleted after incubation through centrifugation. The cells were washed two times with the same buffer to remove the H_2_DCFDA. To cell suspension, different concentrations AgNPs were applied. The fluorescence intensity of DCF by fluorescence spectrophotometer at an excitation wavelength of 488 nm and at an emission wavelength of 535 nm was measured.

### 2.5 Comparative analysis of phytochemical content in extracts and Silver nanoparticles

#### 2.5.1 Determination of Total Phenolic Content

Phenolics are antioxidant molecules with pharmacological activities and reducing capabilities. The total phenolic content in each of the extract and AgNPs synthesized from these extracts were determined quantitatively by using the Folin Ciocalteu method with gallic acid as the standard. Briefly, 20 µL of each extract and AgNPs sample (2 mg/mL) were added to 96-well plate. Then, 90 µL Folin Ciocalteu reagent (1:10 diluted form) was added to the sample. Subsequently 90 µL of 6% sodium carbonate was added. Gallic acid (4 mg/mL stock in methanol) was used as standard positive control. The plate was incubated for 30 mins and then optical density of total phenolic content was measured at 630 nm.

#### 2.5.2 Determination of Total Flavonoid Content

Similar to phenolics, flavonoids possess great medicinal importance and reducing potential. They also show reduction and stabilization potential in the synthesis of AgNPs. The total flavonoid content was measured using Aluminium chloride (AlCl_3_) method. Briefly, 20 µL of each of the extract and AgNPs (2 mg/mL) sample were added to 96-well plate. Potassium acetate (10 µL of 98.15 g/L) was added followed by the addition of 10 µL aluminum chloride (10 g/100 mL). Subsequently, 160 µL of distilled water was added. The 20 µL MeOH instead of sample was used as negative control. The plate was incubated for 30 mins and then the optical density was measured at 405 nm.

#### 2.5.3 DPPH (2,2-diphenylpicrylhydrazyl) assay

A DPPH assay was performed to assess free radical scavenging activity (FRSA) of *S. moorcroftiana* extracts and AgNPs synthesized from it. DPPH is a stable free radical; therefore, it is used to assess whether the extracts and AgNPs possess the ability to scavenge this radical. To determine the antioxidant activity 20 µL of sample (extracts and 2 mg/mL AgNPs) was added to a 96-well plate. Then, 180 µL of DPPH reagent (3.2 mg/100 mL) was added subsequently. Ascorbic acid (4 mg/mL) was used as positive control which is naturally free radical scavenging agent. The plate was incubated for 1 hr and FRSA was measured by recording absorbance at 517 nm.

#### 2.5.4 Oxidation-Reduction potential

The ORP of extracts was determined to know their reduction potential in the synthesis of AgNPs. The ORP of AgNPs was determined to analyze the potential of nanoparticles as oxidative species that are lethal for bacteria. The ORP was measured through ORP sensor probe lubricated with potassium chloride. The rod was dipped in each of the extract and the ORP for each extract sample was measured three times subsequently their average was considered as a final value. Similar procedure was done to determine the ORP of all AgNPs solution (2 mg/ml). The probe was rinsed with distilled water every time it was used for different kind of extract and AgNPs solution.

### 2.6 High-Performance Liquid Chromatography (HPLC) for identification of polyphenols in plant extracts

HPLC quantification was carried out according to a reported method (Zeb 2015). Briefly, 1-g powdered sample of each of the whole and fractionated extract was added to water and methanol in equal ratio, and the mixture was subject to heating in a water bath at 70 °C for 1 h. The mixture was then filtered through a non-pyogenic 0.4 µm CA syringe filter.

To identify and quantify phenolic compounds the Agilent-1260 infinity High-performance liquid chromatography (HPLC) system was used. The HPLC system’s essential parts were a quaternary pump, an auto-sampler, a degasser, and a C18 column (Agilent-Zorbax-Eclipse column). The solution (B and C) gradient was such that solvent B was a mixture of acetic acid: methanol: deionized water (20: 100: 180 v/v), and solvent C was a mixture of acetic acid deionized water: methanol (20: 80: 900) v/v. The solvents were provided as a gradient such that they started and gradually decreased the solvent in concentration. Solvent B was given in volume 100, 85, 50, and 30% at 0, 5, 20, and 25 min, finally giving way to 100% solvent C from 30 min onwards till 40 min. The ultraviolet array detector (UVAD) was set at wavelength 250 nm to analyze phenolic compounds, and the chromatograms were recorded. Phenolic compounds were identified by comparing the retention times of obtained HPLC chromatogram with that of the standards.

## 3. Results and Discussion

### 3.1 Synthesis of silver nanoparticles in plant extracts obtained in solvents with varying polarity

The biosynthesis reaction of AgNPs was performed in each of the extract by allowing it to react with AgNO_3_ solution. After 4 hrs of reaction, a color change was observed that was different for each of the extracts. The change in color is a characteristic determining the reduction of Ag^+^ by plant metabolites (16). This change in color is due to the interaction of light and the surface oscillating electrons of silver nanoparticles, the phenomenon is known as surface plasmon resonance (17). Thus, each of the extract either whole or fraction had the potential of AgNPs synthesis except H-extract. A very slow reaction and thus synthesis of AgNPs were observed in hexane because it is nonpolar and is usually used for the extraction of nonpolar compounds. AgNO_3,_ being polar cannot react with nonpolar compounds. The dried H-extract was dissolved in methanol because of its good extraction capability (18). As a result, H-extract dissolved in methanol when reacted with AgNO_3_ solution, a very low AgNPs synthesis was observed.

### 3.2 UV-Vis spectroscopy of Silver nanoparticles

UV-Vis absorption spectra of the biosynthesized AgNPs showed the characteristic surface plasmon resonance of AgNPs in the range of 400-500 nm, confirming their synthesis (19). The maximum absorbance (represented as λ _(max)_) in this specified range was recorded to be 498 nm for Aq-AgNPs, 432 nm for D-AgNPs, and 414 nm for H-AgNPs (Fig 2 a). (19). Aq-AgNPs and D-AgNPs showed broader peaks indicating the presence of nanoparticles with varying sizes (20). H-AgNPs gave narrow SPR band with little absorption intensity that shows the small concentration of biosynthesized H-AgNPs. (20). With increase in the size of nanoparticles the absorbance intensity decreases (21) as observed with Aq-AgNPs when compared with D-AgNPs. Kochkina and Skobeleva (22) synthesized AgNPs from the reaction of starches dissolved in DMSO with AgNO_3_ and observed the surface plasmon band maximum at 418 nm. The absorption bands of Aq*-AgNPs (λ _(max)_ = 414 nm) and D*-AgNPs (λ _(max)_ = 472 nm) within the characteristic range confirms their presence. The UV-Vis spectra of *S. moorcroftiana* extract in different solvents is taken as control (Fig 2 b) to compare their absorption peaks with that of AgNPs. Although the extracts showed absorbance in the UV-Vis range, no characteristic peaks are found to be similar with the peaks of AgNPs. The D-shaped absorption curve of D*-AgNPs in the range of 400-500 is the indicative of the presence of AgNPs in the sample when compared to D*-extract which absorption showed to be continuously decreasing within this range. Thus UV-Vis spectroscopy results clearly show that the synthesis of AgNPs from their respective extracts is achieved.

**Figure 2:**
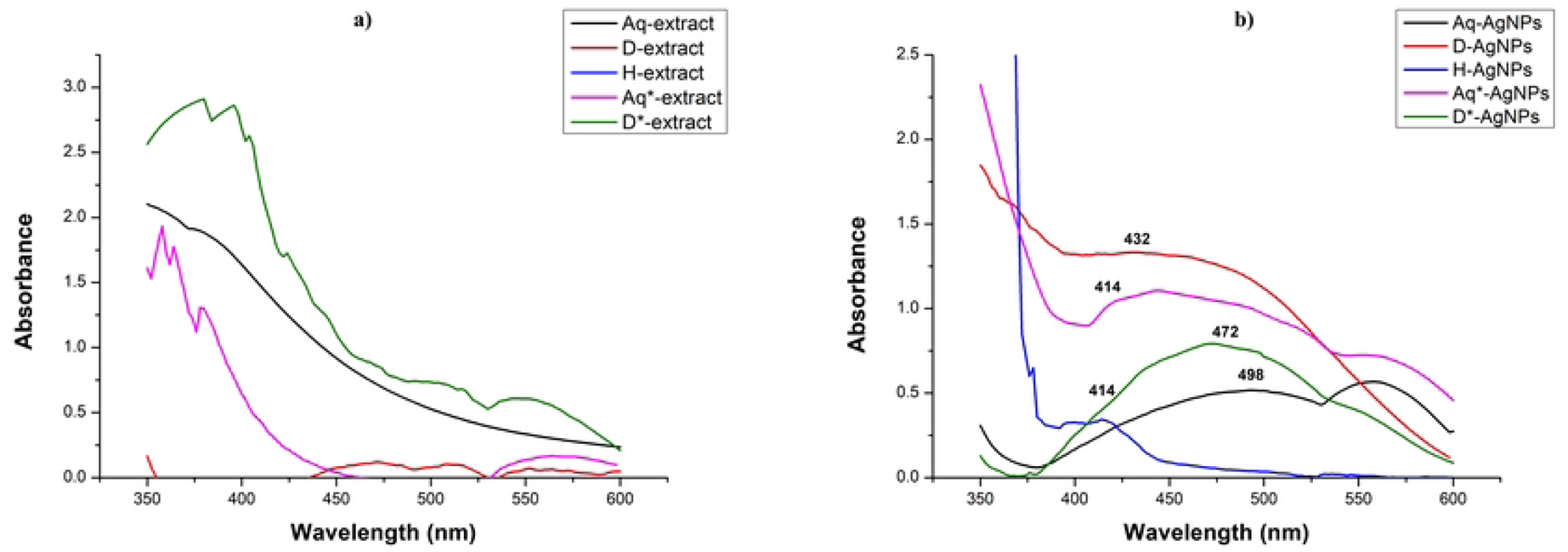
UV-Vis spectra of (a) *S. moorcroftiana extracts* and (b) silver nanoparticles synthesized from these extracts

### 3.3 Fourier Transform Infra-Red Spectroscopy (FTIR)

Generally, the FTIR spectra of the given plant extracts and AgNPs revealed that the compounds present in the extracts have capped the biosynthesized AgNPs and therefore, gave peaks in the same range of wavenumbers. However, less absorption and more transmittance was observed in the FTIR spectra of AgNPs than that of extracts from which they were synthesized.

The FTIR spectra of Aq-extract and Aq-AgNPs (Fig 3a) revealed the absorption of infrared waves by same functional groups within the same wavenumber range. A broad bend was observed in between 3400-3154 cm^-1^ (96% transmittance) for Aq-extract and 3380-3190 cm^-1^ (98% transmittance) for Aq-AgNPs that attributed to the presence of OH stretching (corresponding to alcohol). Also the peaks could be seen at 2980 cm^-1^ (95% transmittance), 2951 cm^-1^ (95% transmittance) and 2972 cm^-1^ (98% transmittance), and 2920 cm^-1^ (97% transmittance) for Aq-extract and Aq-AgNPs, respectively. All these peaks fall in the range which represents the presence of N-H (amine) and C-H (alkane) functional groups, correspondingly. Further, absorption peaks were observed at 1589 cm^-1^, 1060 cm^-1^, and 1034 cm^-1^ (91%, 88%, 88 % transmittance, respectively) for Aq-extract and 1629 cm^-1^, 1060 cm^-1^, and 1055 cm^-1^ (97%, 96%, 96% transmittance, respectively) for Aq-AgNPs. These peaks revealed the occurrence of N-H bending (amine), S=O, and C-O stretching (representing sulfoxide and alcohol groups). A broad peak within the range of 3578-3000 cm^-1^ was observed in the spectrum of D-extract (Fig 3c) along with other noticeable peaks at 1634 cm^-1^, 1047 cm^-1^, and 1019 cm^-1^ Similarly, for D-AgNPs a broad peak was shown within the range of 3500-3100 cm^-1^, and sharp peaks at 1654 cm^-1^, 1615 cm^-1^, and 1030 cm^-1^. The above-mentioned absorption bands attributed to the presence of OH and N-H stretching (corresponding to alcohol and amines), C-H (alkanes), C=C (conjugated alkenes), and C-N (amines). The FTIR spectrum of Aq*-extract (Fig 3b) with in the range of 3524-3040 cm^-1^ showed absorbance (67% transmittance) by showing a broad peak, while in Aq*-AgNPs in the specified range a little absorbance was observed with a slight bend at 3300 cm^-1^ (72% transmittance). This attributed to the presence of OH stretching corresponding to alcohol group. Furthermore, Aq*-extract gave observable peaks at 2941 cm^-1^ (79% transmittance), 1596 cm^-1^ (44% transmittance), 1404 cm^-1^ (54% transmittance), and 1019 cm^-1^ (48% transmittance). The peaks for Aq*-AgNPs were observed at 2920 cm^-1^ (68% transmittance), 1652 cm^-1^ (68 % transmittance), 1400 cm^-1^ (68% transmittance), and 1000 cm^-1^ (60 % transmittance). These peaks represent the presence of OH stretching (alcohol), C-H stretching (alkane), C=C stretching (alkene), S=O stretching (sulfate), and C-F (fluoro compound) respectively, for both extract and AgNPs. A broad peak in spectra of D*-extract and D*-AgNPs (Fig 3c) within the range of 3412-3216 cm^-1^ (66% transmittance) and 3434-3226 cm^-1^ (68% transmittance) representing the presence of OH stretching corresponding to alcohol group. Other noticeable peaks could be seen for D*-extract and AgNPs at 1663 cm^-1^ (76% transmittance) and 1660 cm^-1^ (66% transmittance), 1178 cm^-1^ (50% transmittance) and 1168 cm^-1^ (62% transmittance), 1045 cm^-1^ (36% transmittance) and 1027 cm^-1^ (54% transmittance). The absorption at these wavenumbers showed the presence of OH stretching (alcohol), C=C (alkene), C-O stretching (ester), and C-N stretching (amine), respectively.

**Figure 3:**
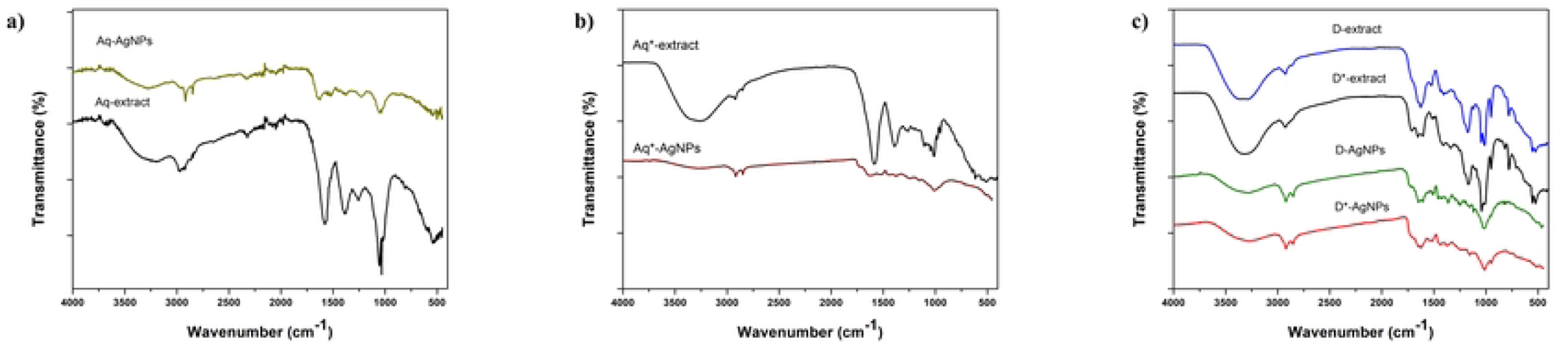
FTIR spectra of a) Aq-extract and AgNPs b) Aq*-extract and AgNPs c) D-extract and AgNPs, D*-extract and AgNPs

The presence of these functional groups indicates the presence of polyphenols (such as flavonoids and phenolics), terpenoids, carboxylic acid, amines in proteins, aromatic amines, alkenes, alkanes, and organosulfure compounds attached to the surface of AgNPs. The presence of these phytochemicals in both the plant extracts and AgNPs indicate their role in the synthesis and stabilization of AgNPs (23–25). It is challenging to point out a single specific class of phytochemicals involved in the synthesis of AgNPs, but generally, all these compounds are significant for the synthesis of silver nanoparticles (26). These phytochemicals possess antibacterial properties and can result in the synthesis of AgNPs with enhanced antibacterial activity (27). Only D*-extract and D*-AgNPs showed a distinctive peak at 1168 cm^-1^ and 1178 cm^-1^ indicating C-O vibration corresponding to esters. In previous studies it has been shown that ester molecules were obtained in high concentration through extraction in polar solvents as compared to nonpolar (28). Moreover, both aqueous whole and fraction extracts and AgNPs based on them revealed the presence of sulfate or sulfoixde (S=O) that is the functional group of organosulfur compounds in plants (29). This compound was not found in DMSO based extracts and AgNPs because they are known to decompose in DMSO (30).

### 3.4 Transmission electron microscopy of silver nanoparticles

TEM was used to analyze the morphology of biosynthesized AgNPs. The TEM images confirmed the synthesis of AgNPs in all the extracts of *S. moorcroftiana* (Fig 4). The nanoparticles in all the extracts appeared to be poly dispersed having both large and small diameter with average diameter of 17.889, 17.11, 18.82, and 17.815 nm for Aq-AgNPs, D-AgNPs, Aq*-AgNPs, and D*-AgNPs, respectively (Table 1). All of the biosynthesized AgNPs were almost spherical in shape as determined by TEM micrographs. AgNPs with smaller sizes and spherical morphology are more effective against bacteria because they can easily attach and enter the cells via membranes leading to bactericidal effects (4). The close analysis of images showed that clusters of nanoparticles are surrounded by a layer. Such layers are found around nanoparticles mainly synthesized from medicinal plant extracts, where the phytochemicals form capping layer and help to shape the particles during growth (31).

**Figure 4:**
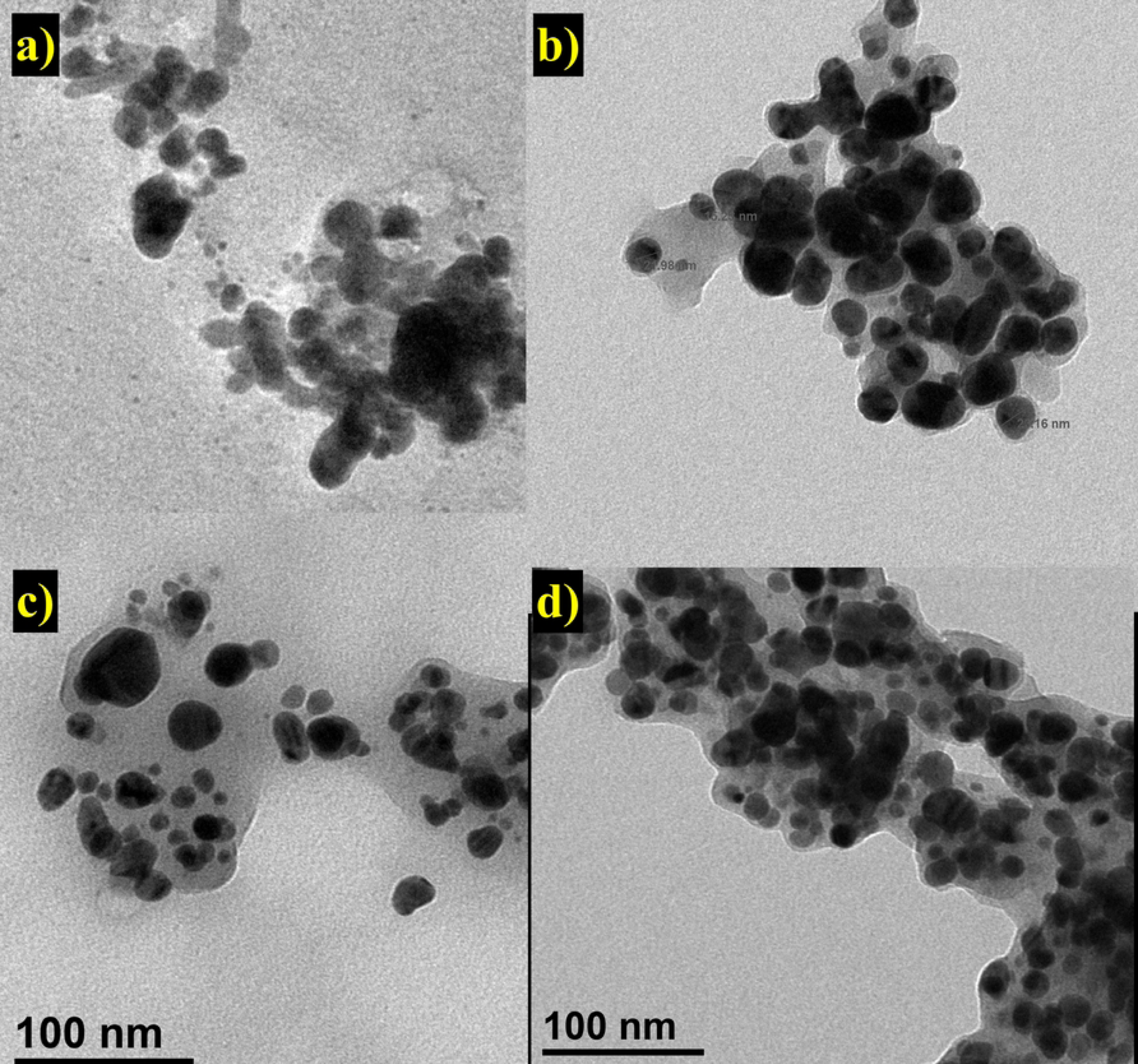
Transmission electron microscopy image of a) Aq-AgNPs, b) Aq*-AgNPs, c) D-AgNPs and d) D*-AgNPs

**Table 1:** Combined Table of size and stability-based characteristics of Silver nanoparticles

| Types of AgNPs | PDI | Hydrodynamic Size (nm) | Zeta potential (mV) | TEM (nm) |
| --- | --- | --- | --- | --- |
| D*-AgNPs | 0.335 | 377 | -21.8±0.16 | 17.815 |
| D-AgNPs | 0.266 | 365 | -18.5±0.24 | 17.11 |
| Aq*-AgNPs | 0.5 | 99.36 | -16.3±0.26 | 18.82 |
| Aq-AgNPs | 0.373 | 135.7 | -17.6±0.14 | 17.889 |

### 3.5 Zeta potential data for assessing the stability of silver nanoparticles

Zeta potential, the electrostatic repulsion or attraction between the particles is determined through Dynamic Light Scattering (DLS). The negative or positive zeta values indicate the repulsive force between the particles showing dispersity and stability. All types of samples showed negative zeta potential values such as −17.6±0.14 mV, −18.5±0.24 mV, −16.3±0.26 mV, and −21.8±0.16 mV for Aq-AgNPs, D-AgNPs, Aq*-AgNPs, and D*-AgNPs, respectively (Fig 5). The negative zeta potential values indicated that the AgNPs possessed a positive charge in solution that attracted negative charges to surround them (32). The negative zeta values also indicate the repulsive force among the nanoparticles, hence confirms their stability (33). In this context, the DMSO plant extract based AgNPs were found to be more stable than the aqueous one. Furthermore, between the DMSO plant extract based AgNPs, D*-AgNPs revealed to be more stable with zeta value = −21.8±0.16 mV.

**Figure 5:**
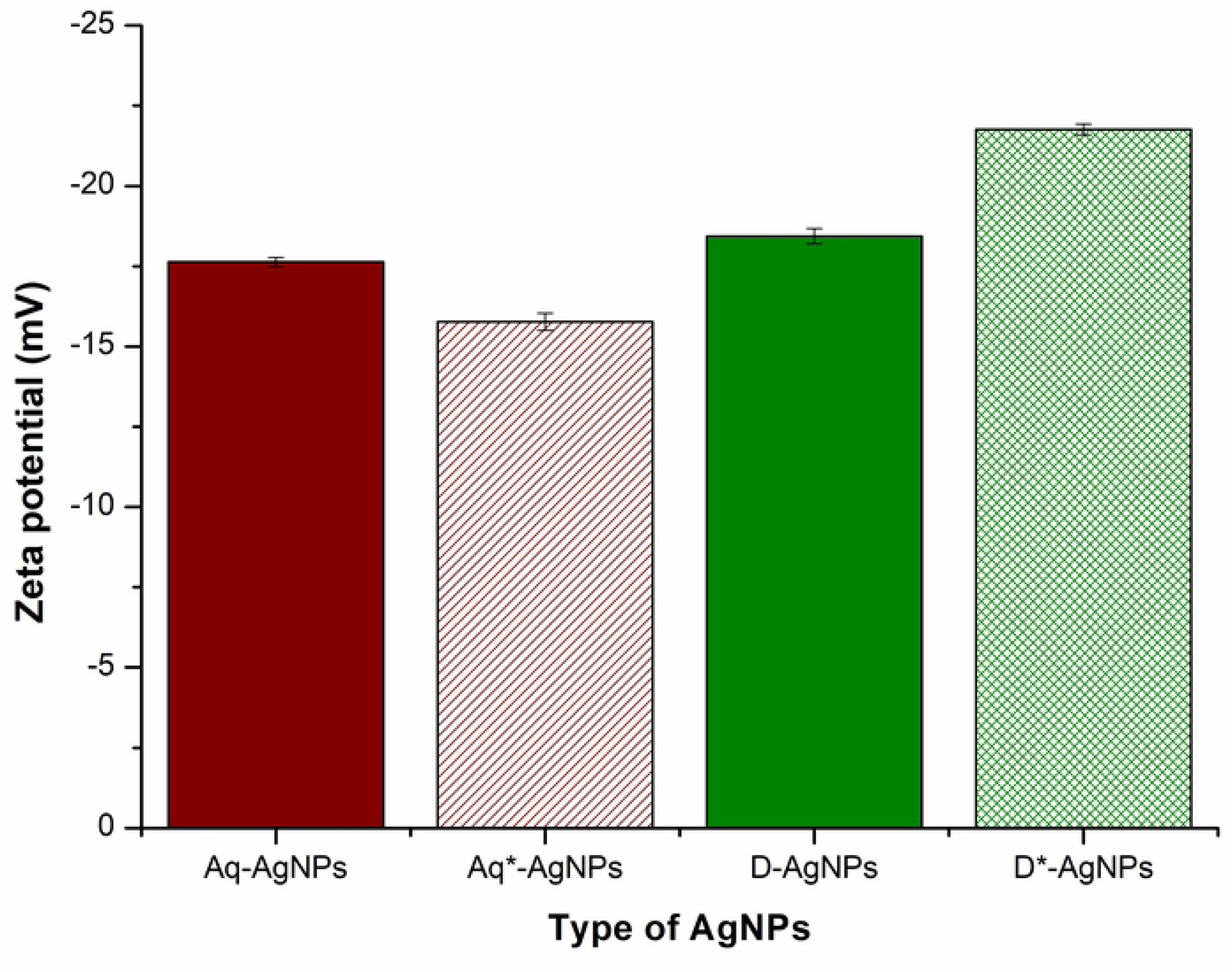
Determination of Zeta potential of silver nanoparticles synthesized via different solvents

### 3.6 Size distribution of silver nanoparticles determined through Dynamic light scattering

DLS was used to determine the polydispersity index, referring to the size distribution of nanoparticles in solution. Furthermore, hydrodynamic size or average zeta size was also calculated for four types of AgNPs i.e. Aq-AgNPs, D-AgNPs, Aq*-AgNPs, and D*-AgNPs (Fig 6). The values of the mentioned characteristics are shown in Table 1. The results revealed the average zeta size of Aq-AgNPs, D-AgNPs, Aq*-AgNPs, and D*-AgNPs to be 135.7 d. nm, 365 d. nm, 99.36 d. nm, and 377 d. nm, respectively. All the AgNPs were found to be less polydispersed with highest PDI being 0.5 recorded for Aq*-AgNPs followed by Aq-AgNPs with 0.373 PDI, D*-AgNPs with 0.335 PDI, and D-AgNPs with the lowest PDI (0.26). The size determined by DLS was found to be larger than the size determined by TEM because it measures hydrodynamic size of nanoparticles rather than their physical diameters (34). This means that during movement of nanoparticles in liquid medium, the electric dipole layer of the solvent surrounds them which can also influence their motion in the solvent. Thus, hydrodynamic diameter shows the combined diameter of metal core and the solvent molecules surrounding it (35). In this context, DMSO based AgNPs are probably surrounded by more protective layers than aqueous extract based AgNPs, thus having large hydrodynamic diameter.

**Figure 6:**
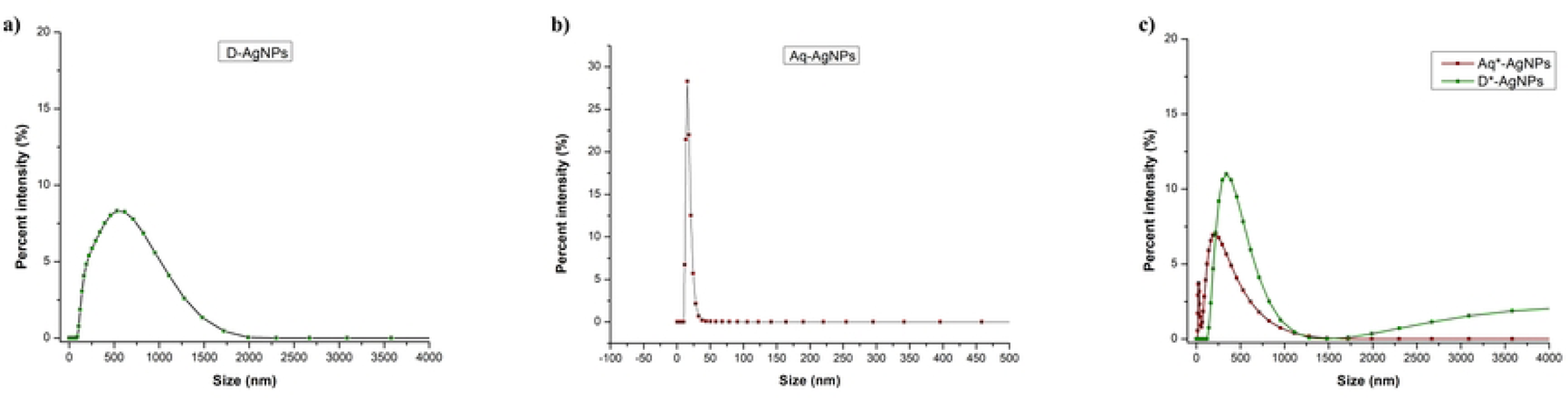
Size distributions of silver nanoparticles synthesized via different extracts

### 3.7 Comparative analysis of phytochemical content in plant extracts and Silver nanoparticles

#### 3.7.1 Total phenolic content in extracts and silver nanoparticles

The TPC and TFC results showed that high amount of phenolic and flavonoid compounds are present in the extracts which act as reducing and capping agents utilized in the synthesis of stable AgNPs (36). The TPC and TFC of AgNPs gave evidence of the capping and stabilizing capability of these compounds (37) showing that during synthesis, they attach to the AgNPs to stabilize them and provide them enhanced pharmacological properties.

TPC of extracts showed diverse results such as 98.92±4.12, 255.50±3.13, 262.48±0.84, 227.76±0.46, and 67.42±1.21 µg GAE/mL for Aq-extract, Aq*-extract, D-extract, D*-extract, and H-extract, respectively (Fig 7). The highest phenolic content was found in D-extract, followed by Aq*-extract. The least phenolics were extracted by hexane. The TPC of AgNPs was substantially lower than that of extracts (Fig 7). 13.01±1.19, 13.01±1.19, 26.8±1.95, and 23.44±0.19 µg GAE/mL are the concentrations of phenolics bound to Aq-AgNPs, Aq*-AgNPs, D-AgNPs, and D*-AgNPs, respectively. Compared to Aqueous plant extract-based AgNPs, D-AgNPs and D*-AgNPs had increased levels of phenolic compounds.

**Figure 7:**
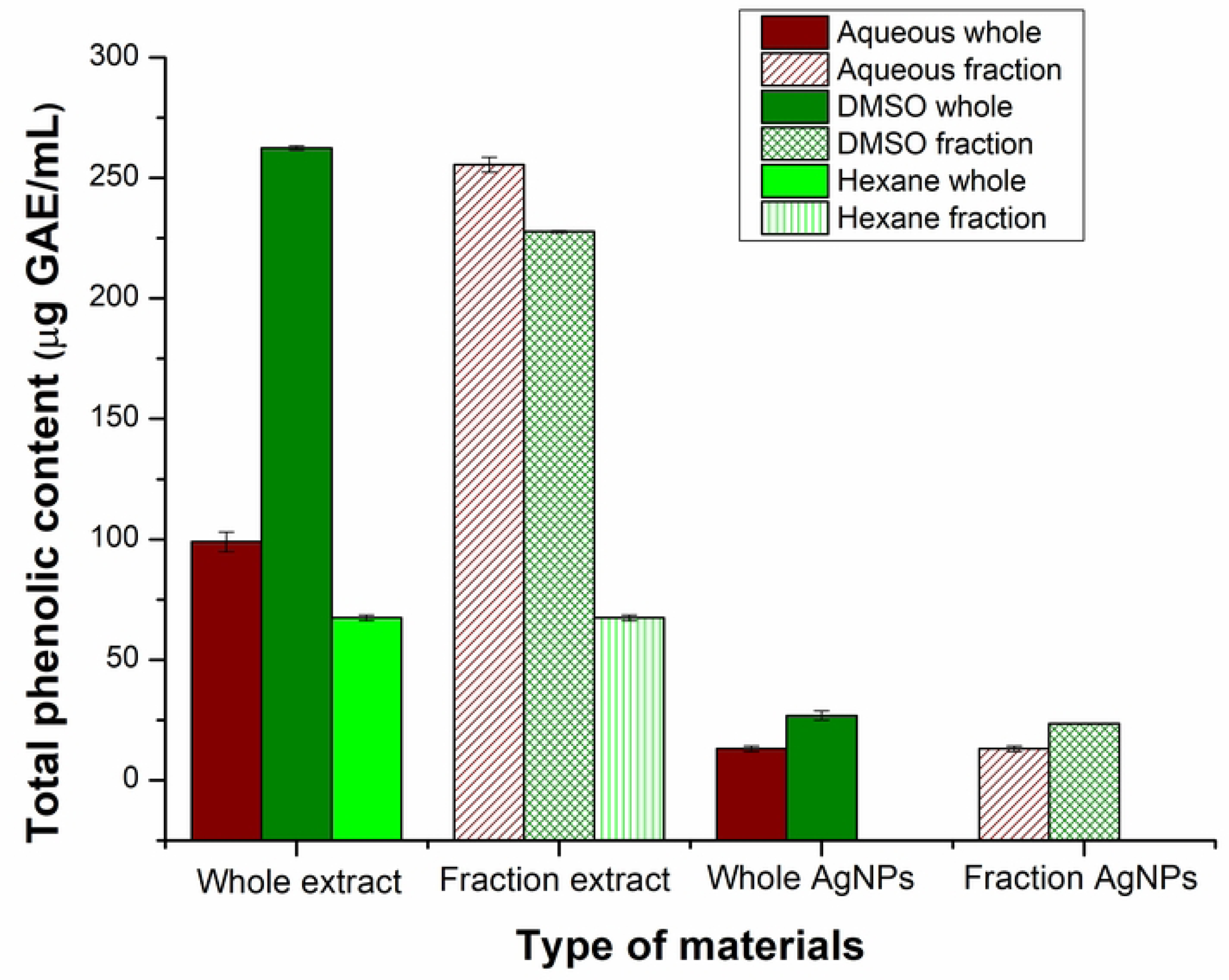
Comparative analysis of total phenolic content in different extracts and AgNPs based on different solvent-based extracts. Hexane based AgNPs synthesis was negligible and could not be analyzed for TPC

#### 3.7.2 Total flavonoid content of extracts and silver nanoparticles

The flavonoids were present in higher quantity in extracts than AgNPs. The D-extract and D*-extract had the same and highest flavonoid content i.e., 11.42±0.54 µg QE/mL, and 11.42±0.3 µg QE/mL, respectively. TFC of Aq-extract, Aq*-extract, and H-extract was calculated to be 7.06±2.29 µg QE/mL, 5.40±0.61 µg QE/mL, and 4.72±0.13 µg QE/mL, respectively. 3.58±0.29 µg QE/mL, 9.2±0.25 µg QE/mL, 6.17±0.11 µg QE/mL, and 5.02±0.19 µg QE/mL TFC was recorded for Aq-AgNPs, Aq*-AgNPs, D-AgNPs, and D*-AgNPs. Like TPC, D and D*-AgNPs gave high TFC value (Fig 8).

**Figure 8:**
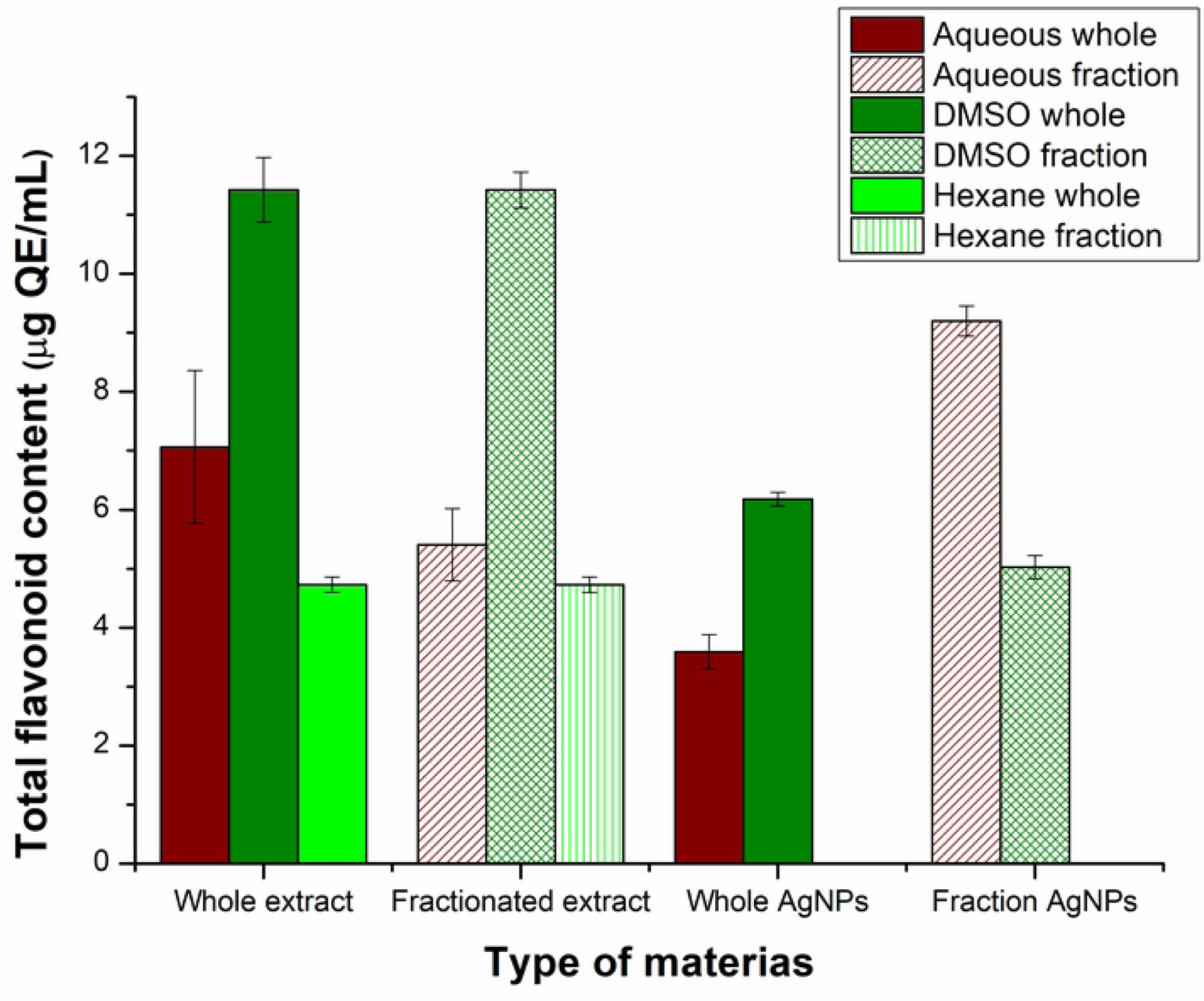
Comparative analysis of total flavonoid content in extracts and silver nanoparticles based on different solvent-based extracts.

#### 3.7.3 Antioxidant activity of extracts and silver nanoparticles

The antioxidant activity of extracts and AgNPs was determined by calculating their free radical scavenging activity (FRSA) (Fig 9). This free radical scavenging property is the result of reductants having reducing power that can react with and prevent the formation of free radicals (38). The results of free radical scavenging activity indicate effective antioxidant activities of all types of extracts. This could be linked with the presence of TPC and TFC, because, phenolic compounds possess strong antioxidant potential that are found in considerable amounts in plant extracts (39). Among the extracts, D*-extract showed the highest percent of free radical scavenging activity i.e., 80.8±0.72%, which can be attributed to the increased quantity of phenols and flavonoids extracted by DMSO in fraction. DMSO is a preferred solvent for the extraction of phenolic and flavonoid compounds as in both cases DMSO based plant extracts revealed their higher concentration. Demir, Turan (40) used DMSO for extraction of *Rhododendron luteum* and revealed its high antioxidant activity due to the extraction of higher quantities of polyphenols. This free radical scavenging property is the result of reductants having reducing power that can react with and prevent the formation of free radicals (38). The H-extract also revealed good antioxidant activity having 74.36±1.0 percent free radical scavenging activity.

**Figure 9:**
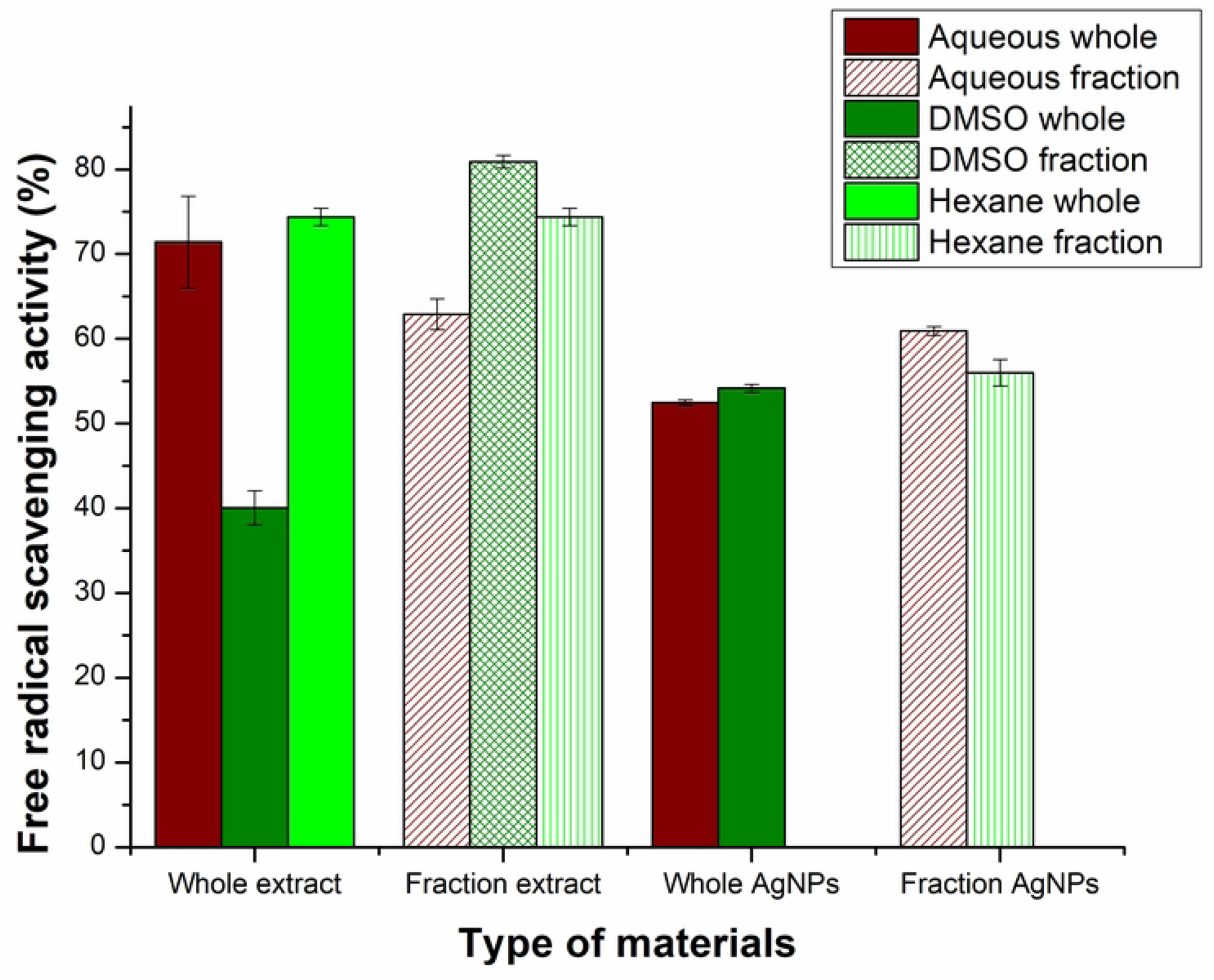
Antioxidant activity of plant extracts and silver nanoparticles synthesized from these extracts

AgNPs synthesized from these extracts were also found to be good antioxidant agents with the highest antioxidant potential (61±0.50%) of Aq*-AgNPs followed by D*-AgNPs with 56±1.55% FRSA. This reveals that AgNPs synthesized from fractionated plant extracts was higher as compared to the one synthesized from whole extracts. Thus, the strategy of fractionated plant extraction and the synthesis of AgNPs from these extracts result in the production of good antioxidant agents.

#### 3.7.4 Oxidation Reduction potential of extracts and silver nanoparticles

ORP of extracts was determined to find the reducing agents and their reduction potential for the synthesis of AgNPs. The more negative is the value the more reducing agents are present in the extract having the capability to reduce Ag^+^ to AgNPs indicating high reduction potential (41). The highest reduction potential was recorded for Aq-extract (−113.667±1.17 mV) and the lowest for H-extract (−3.66±0.69 mV) while Aq*-extract gave positive ORP value (67.33±0.19 mV). The given ORP values of AgNPs show their high oxidation potential and hence bactericidal properties. Higher positive ORP values indicate high concentration of oxidizing agents present in a solution.

**Table 2:** Comparative analysis of Oxidation Reduction Potential of extracts and AgNPs synthesized from these extracts

| ORP of extracts (mV) |  |  |  |  |
| --- | --- | --- | --- | --- |
| Aq-extract | Aq*-extract | D-extract | D*-extract | H-extract |
| $-113.66 \pm 2$ | $67.33 \pm 0.33$ | $-4.66 \pm 2.66$ | $-65.33 \pm 1.45$ | $-3.66 \pm 1.2$ |
| ORP of AgNPs (mV) |  |  |  |  |
| Aq-AgNPs | Aq*-AgNPs | D-AgNPs | D*-AgNPs | H-AgNPs |
| $44 \pm 0.57$ | $110 \pm 3$ | $70.33 \pm 2.9$ | $92.33 \pm 4.25$ | $-110 \pm 1.66$ |

### 3.8 Identification of polyphenols through HPLC

The results of HPLC revealed the whole and fractionated extracts of S. moorcroftiana to be enriched with phenolic acids and flavonols. Such composition of the extracts make them efficient source for the synthesis of AgNPs. From the chromatogram of each of the extract, those with higher peak area (%) are picked for analysis (Table 3). Polyphenols found in Aq-extract include malic acid (25.49), caftaric acid (8.387), and myricetin (32.04). D-extract contains gallic acid (14.746), rosmarinic acid (46.227), and vitamin C (4.555). Aq*-extract possess rutin (23.884), Syringic acid (15.57), Quercitin 3,7-di-o-glucoside (47.30), and Kaempferol-3-(P-coumoryl-diglucoside)-7-glucoside (7.867) while D*-extract possess Kaempferol-3-(P-coumaroyl-diglucoside)-7-glucoside (7.867), Quercetin-3-d-galactoside (5.685), and Caffeic acid (83.03). H-extract contained the polyphenols like Gallocatechin (52.89), Gallic acid (3.230), and Kaempferol-3-feryloyl sophoroside-7-glucoside (16.00).

**Table 3:** Polyphenolic compounds with their retention time and peak area (%)

| Sample extract | Retention time | Compounds identity | Peak area (%) | Identification reference |
| --- | --- | --- | --- | --- |
| Aq-extract | 2.326 | Malic acid | 25.49 | Standard |
|  | 3.209 | Caftaric acid | 8.387 | Standard |
|  | 3.775 | Gentisic acid | 0.173 | Standard |
|  | 24.814 | Quercitin-7-o-sophoroside | 0.282 | Standard |
|  | 26.368 | Quercitin dihydrate | 0.415 | Standard |
|  | 28.662 | Pyrogallol | 1.384 | Standard |
|  | 31.152 | Quercitin-3-(caffeoyl diglucoside)-7-glucoside | 0.135 | Standard |
|  | 34.246 | Rosmarinic acid | 1.222 | Standard |
|  | 34.902 | myricetin | 32.04 | Standard |
| D-extract | 4.349 | Gallic acid | 14.746 | Standard |
|  | 4.868 | Vitamin C | 46.227 | Standard |
|  | 34.249 | Rosmarinic acid | 4.555 | Standard |
| Aq*-extract | 12.832 | Morin | 0.086 | Standard |
|  | 14.541 | P-coumaric acid | 0.0717 | Standard |
|  | 20.309 | Quercitin-3-(caffeoyl diglucoside)-7-glucoside | 2.987 | Standard |
|  | 22.179 | Syringic acid | 15.57 | Standard |
|  | 22.835 | Rutin | 23.884 | Standard |
|  | 23.892 | Quercitin 3,7-di-o-glucoside | 47.30 | Standard |
|  | 25.439 | Kaempferol 3-(caffeoyl-diglucoside)-7-glucoside | 6.779 | Standard |
|  | 35.418 | Kaempferol-3-feryloyl sophoroside-7-glucoside | 2.368 | Standard |
| D*-extract | 18.145 | Kaempferol-3-(P-coumoryl-diglucoside)-7-glucoside | 7.867 | Standard |
|  | 29.621 | Caffeic acid | 83.03 | Standard |
|  | 30.44 | Mandelic acid | 0.337 | Standard |
|  | 34.311 | Rosmarinic acid | 0.392 | Standard |
|  | 35.741 | Kaempferol-3-feryloyl sophoroside-7-glucoside | 0.125 | Standard |
| H-extract | 5.51 | Gallocatechin | 52.89 | Standard |
|  | 4.25 | Gallic acid | 3.230 | Standard |
|  | 6.358 | Caffeic acid | 0.309 | Standard |
|  | 35.663 | Kaempferol-3-feryloyl sophoroside-7-glucoside | 16.00 | Standard |

Plants being the natural reducing agents contain different secondary metabolites in the form of condensed polyphenols which takes part in the reduction of metal salts and stabilization of their nano forms. Identification of these secondary metabolites is vital to understand the possible reaction between the precursor and extract used (42). Various types of polyphenols are identified in each of the extract of *S. moorcroftina.* With the help of the reaction of these phenols and flavonoids with AgNO_3_, stable AgNPs in high concentration were produced. These polyphenols have been reported to possess strong antioxidant and antibacterial activity. Among the extracts, only to the D*-extract, all tested strains of bacteria were susceptible. This can also be linked to the results of TPC and TFC that show D*-extract possessing higher TFC and TPC values. Despite the presence of polyphenols in Aq and Aq*-extract, none of them showed any antibacterial activity. This is similar to the results of a study where the ethanolic extract of *Crataegus monogyna* showed good activity against Gram positive and negative bacteria while its aqueous extract did not show any. The antibacterial activity was attributed to the presence of high polyphenol content (43). Caffeic acid is a known potent antibacterial agent which in the present study is found in high amount in D*-extract (44). In addition, kaempferol and its derivatives have been shown in previous studies to possess good antibacterial potential against Gram positive and negative bacteria (45). Thus, the fractionated DMSO based extract of *S. moorcroftiana* was able to extract polyphenols that were biologically active against pathogenic bacteria.

### 3.9 Antibacterial activity of extracts and silver nanoparticles

#### 3.9.1 Disk diffusion assay of extracts and silver nanoparticles

The antibacterial activity of whole and fractionated *S. moorcroftiana* extracts and AgNPs synthesized from these extracts was examined. All types of AgNPs showed moderate activity against multi drug resistant human pathogens such as *Klebsiella pneumoniae, Pseudomonas aeruginosa, Staphylococcus aureus,* and *Enterococcus faecalis* (Table 4). Among the extracts, D*-extract showed good antibacterial potential with inhibition zones having diameter of 11 mm, 9.83 mm, 9.83 mm, and 10.33 mm against *K. pneumoniae, P. aeruginosa, S. aureus, and E. faecalis,* respectivley. The extraction of phytochemicals depends on method of extraction, solvent, and time duration. Here, solvent is considered as a main factor. On this basis, only DMSO based fraction of *S. moorcroftiana* rather than its aqueous extracts was able to show antibacterial effect against all the tested strains. This is probably because the plant components active against bacteria may not be effectively extracted in water while the antibacterial aromatic or saturated organic compounds are mostly obtained through organic solvent extraction (46). As compared to D-extract, D*-extract had higher antibacterial potential. When the antibacterial activity of D*-extract is linked to the results of FTIR they indicate the presence of esters that might be responsible for its antibacterial activity because esters have been reported to possess good antibacterial activity (47). D*-extract also possessed concentrated biological important phytochemicals such as polyphenols and flavonoids. Kumar, Nehra (48) showed that *E. faecalis* is most sensitive to plant compounds such as polyphenols. Therefore, this specific strain (as compared to other strains) in our study showed increased sensitivity to H-extract, D-extract, and D*-extract.

**Table 4:** Inhibition zone sizes of AgNPs against MDR bacteria

| Bacterial strains | Zone of inhibition (mm) |  |  |  |  |  |  |  |  |  |
| --- | --- | --- | --- | --- | --- | --- | --- | --- | --- | --- |
|  | H-AgNPs | Aq*-AgNPs | D*-AgNPs | Aq-AgNPs | D-AgNPs | H-extract | Aq*-extract | D*-extract | Aq-extract | D-extract |
| *Klebsiella pneumonia* | 8±0 | 11±0 | 11±0 | 12±0 | 11±0 | 0 | 0 | 11±0 | 0 | 0 |
| *Pseudomonas aeruginosa* | 9.33±0.33 | 10.33±0.33 | 12 | 11.66±0.33 | 9.33±0.33 | 0 | 0 | 9.83±0.44 | 0 | 0 |
| *Staphylococcus aureus* | 8.67±0.33 | 10.66±0.33 | 10.5±0.28 | 11±0 | 9.83±0.16 | 0 | 0 | 9.83±0.44 | 0 | 0 |
| *Enterococcus faecalis* | 8.60±0.16 | 10±0 | 11.33±0.33 | 11.5±0.28 | 11±0.57 | 11.3±0 | 0 | 10.33±0.33 | 0 | 10±0 |

The largest zone of inhibition was 12 mm that was recorded for Aq-AgNPs against *K. pneumoniae* and D*-AgNPs against *P. aeruginosa.* When compared, Aq-AgNPs gave larger zone of inhibition than Aq*-AgNPs but in case of DMSO extract based AgNPs, D*-AgNPs showed higher antibacterial activity than D-AgNPs. H-AgNPs gave comparatively smaller zones of inhibition i.e. 8 mm, 9.33 mm, 8.67 mm, and 8.60 mm against *K. pneumoniae, P. aeruginosa, S. aureus, and E. faecalis,* respectivley, revealing it to be weak antibacterial agent. Four types of controls were taken alongside i.e. dH_2_O, DMSO, hexane, and ceftadizime antibiotic (200µg/ml).

Each type of AgNPs showed activity against all tested MDR strains. H-AgNPs was the least effective because H-extract did not contain enough phytochemicals to synthesize and cap AgNPs. As in our results we can see that D*-AgNPs are more stable, have higher antioxidant activity, contain increased amount of phenolic and flavonoid compounds, and also have the high oxidation potential which should how make them most effective against resistant bacteria. However, Aq-AgNPs were the most effective with large inhibition zone against each bacterium as compared to all other AgNPs. The antibacterial results linked with FTIR analysis revealed the presence of S=O (sulfoxide) on Aq-AgNPs that is commonly found in organosulfur compounds having strong antibacterial activity along with other phytochemicals (49). For instance, thiosulfinate (organosulfur compound) such as allicin has been recognized as responsible for the antibacterial property of garlic and onion (50). The contributing role of these compounds with other phytochemicals may be the reason for enhanced antibacterial activity of Aq-AgNPs. Though the same group was also found in Aq*-AgNPs, their antibacterial potential was not that effective because at the end of fractionation (solvent-solvent extraction), water had few remaining phytochemicals to extract and further utilize for synthesis and capping of AgNPs. Such organosulfur compounds are immiscible in water and are often more soluble in less polar solvents than water (30, 51). Therefore, these compounds were not biologically available in Aq-extract and Aq*-extract for interaction with bacteria, hence did not show any antibacterial activity.

#### 3.9.2 Minimum Inhibitory concentration (MIC) of AgNPs

The minimum inhibitory concentrations of AgNPs against all aforementioed bacterial strains were determined through microdilution assay. The bacterial growth inhibition with the application of AgNPs was determined through visual observation of turbidity in the wells of microtitre plate. Each kind of AgNPs showed different MIC values against each bacterial strain as shown in table 5. The lowest MIC of various AgNPs against the given bacterial strains are: 7.8 µg/mL of H-AgNPs against *P. aeruginosa,* 31.25 µg/mL of Aq*-AgNPs against *P. aeruginosa* and *Staphylococcus aureus,* 31.25 µg/mL of D*-AgNPs against *P. aeruginosa* and *E. faecalis,* 7.8 µg/mL of Aq-AgNPs against *P. aeruginosa,* and 31.25 µg/mL of D-AgNPs against *P. aeruginosa*. Thus, from these results, it is clear that *P. aeruginosa* is the most sensitive to all kinds of AgNPs even at very low concentrations. Similar results were shown by de Lacerda Coriolano, de Souza (52) where *P. aeruginosa* was sensitive to the concentrations of AgNPs between 1 and 200 µg/mL. All types of AgNPs were effective in their lower concentrations that ultimately reduce their cytotoxic effects. However, like the results of disk diffusion assay, Aq-AgNPs against each bacterium compared to other AgNPs revealed to be relatively effective in lower concentrations such as 15.625 µg/mL, 7.8 µg/mL, 31.25 µg/mL, and 15.625 µg/mL against *K. pneumoniae, P. aeruginosa, S. aureus,* and *E. faecalis*, respectively.

**Table 5:** Minimum inhibitory concentration of AgNPs against resistant pathogenic strain

| Bacterial strains | Minimum inhibitory concentration (µg/mL) |  |  |  |  |
| --- | --- | --- | --- | --- | --- |
|  | H-AgNPs | Aq*-AgNPs | D*-AgNPs | Aq-AgNPs | D-AgNPs |
| *Klebsiella pneumonia* | 250 | 125 | 62.5 | 15.625 | 62.5 |
| *Pseudomonas aeruginosa* | 7.8 | 31.25 | 31.25 | 7.8 | 31.25 |
| *Staphylococcus aureus* | 500 | 31.25 | 62.5 | 31.25 | 125 |
| *Enterococcus faecalis* | 500 | 250 | 31.25 | 15.625 | 62.5 |

#### 3.9.3. Demonstration of the mechanism of action of bacterial inhibition by AgNPs

One of the main mechanisms through which antimicrobials induce bactericidal effect is the generation of reactive oxygen species. These ROS targets and alter the sites of amino acids and nucleotides, resulting in protein and DNA damage, causing bacterial cell death (53). The results of ROS quantification shows the production of ROS in all three types of bacteria by all types of applied AgNPs (Fig 10). The highest level of ROS was generated varyingly in each of the bacterial culture, such as, Aq*-AgNPs was able to produce increased level of ROS in *K. pneumonia* and *S. aureus*, while in *P. aeruginosa,* D*-AgNPs produced increased level of ROS. Also, in each bacterial culture, Aq-AgNPs consistently generated significant amount of ROS. A similar trend with respect to the relation of concentration of AgNPs and fluorescence intensity of ROS was noted in *K. pneumoniae* and *P. aeruginosa*. In these bacterial cultures at low concentrations of AgNPs (31.2-62.5 µg/mL), the ROS are generated in high numbers, but as soon as the concentration of AgNPs increases (reaches 250 µg/mL), the number of ROS falls gradually. This might occur because with the application of higher concentrations of AgNPs the bacterial cells die and ultimately they are not able to produce ROS (54). However, in *S. aureus,* the number of ROS retained to increase with increase in the concentration of AgNPs. Thus, all AgNPs were shown to be applicable as antibacterial agents because of their reasonably high inhibitive activity even in very less concentrations.

**Figure 10.**
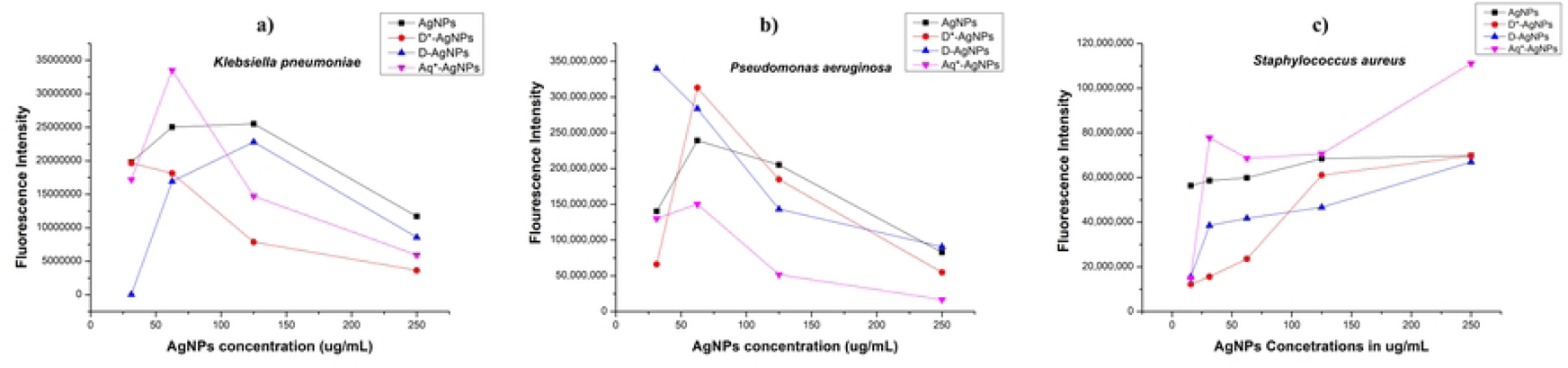
Reactive oxygen quantification in bacterial cultures grown in the presence of different concentrations of different types of AgNPs

## 4. Conclusion

Green synthesized AgNPs because of their ecofriendly synthesis and enhanced antibacterial activity have gained enormous attention of researchers to provide solution for antibiotic resistance. In this study, the extract of *S. moorcroftiana* (whole and fraction) was used for the synthesis of AgNPs in three solvents (water, DMSO, and hexane) with decreasing polarity. The characterization techniques confirmed the synthesis of spherical AgNPs in high concentration with average diameter less than 20 nm, having high polydispersity and stability. Phytochemicals like proteins, flavonoids, phenolic compounds and organic acids are identified to be significantly involved in the synthesis of AgNPs. These compounds, when extracted in fractions show their targeted role in the synthesis of AgNPs and their antibacterial effect, as observed in D*-extract. The plant compounds extracted in DMSO after extraction through hexane showed good reducing, stabilizing, and antibacterial potential. Regardless of identifying the targeted role of phytochemicals, Aq-AgNPs were found to be most effective antibacterial agents because of diverse phytochemicals (wholly extracted in water) capped on them. Overall, this study showed that the targeted extraction of phytochemicals from medicinal plant like *S. moorcroftiana* can help to obtain the antimicrobial plant components that can be applied against bacteria and used for the synthesis of AgNPs with enhanced antibacterial activity and reduced toxicity.

## Acknowledgements

The authors acknowledge the role of the University of Malakand in providing a conducive environment for research activities. The authors acknowledge the role of the Centralized Resource Laboratory, University of Peshawar for providing analytical facilities. The authors also acknowledge the role of the international foundation for Science in Sweden. Tauqir A. Sherazi acknowledges the financial support from the Alexander von Humboldt Foundation.

## Author contributions

**Maham Khan:** Methodology, Data curation, Writing-Original draft preparation; **Tariq Khan:** Conceptualization, Supervision, Reviewing and Editing; **Muhammad Aasim:** Software, Validation; **Tauqeer A. Shirazi:** Analysis, Software; **Shahid Wahab:** Analysis, Visualization; **Muhammad Zahoor:** Investigation, Interpretation.

## Conflict of Interest

The authors declare no conflict of interest.

## Notes

### Competing Interest Statement

The authors have declared no competing interest.

